# Temperature-mediated inhibition of a bumble bee parasite by an intestinal symbiont

**DOI:** 10.1101/360263

**Authors:** Evan C Palmer-Young, Thomas R Raffel, Quinn S McFrederick

## Abstract

Competition between organisms is often mediated by environmental factors including temperature. In animal intestines, nonpathogenic symbionts compete physically and chemically against pathogens, with consequences for host infection. We used metabolic theory-based models to characterize differential responses to temperature of a bacterial symbiont and a co-occurring trypanosomatid parasite of bumble bees, which regulate body temperature during flight and incubation. We hypothesized that inhibition of parasites by bacterial symbionts would increase with temperature, due to symbionts having higher optimal growth temperatures than parasites.

We found that a temperature increase over the range measured in bumble bee colonies would favor symbionts over parasites. As predicted by our hypothesis, symbionts reduced the optimal growth temperature for parasites, both in direct competition and when parasites were exposed to symbiont spent medium. Inhibitory effects of the symbiont increased with temperature, reflecting accelerated growth and acid production by symbionts. Our results indicate that high temperatures, whether due to host endothermy or environmental factors, can enhance the inhibitory effects of symbionts on parasites. Temperature-modulated manipulation of microbiota could be one explanation for fever- and heat-induced reductions of infection in animals, with consequences for diseases of medical and conservation concern.

## INTRODUCTION

Temperature governs rates of the chemical interactions that underlie life, growth, and reproduction, shaping biological processes from the level of the enzyme to the ecosystem [1]. One area of biology where temperature has demonstrated effects is on species interactions such as parasitism, where temperature can have profound effects on infection outcomes and transmission[2,3]. High host body temperatures have been shown to reduce infection intensity and infection-related mortality in both plants and animals [4–6], and metabolic and behavioral fevers are common responses to infection in vertebrates and insects [4,7,8].

Another factor that can influence infection outcome is the host-associated microbiota. The microbiota of the skin and gut constitute barriers to infection that can physically and chemically interfere with pathogen invasion, as well as modify host immune responses [9]. Because microbial taxa can differ widely in their optimal growth temperatures, alterations in temperature can affect the relative competitive abilities of co-occurring species [10]. These differential responses of interacting species to temperature, referred to variously as “asymmetries” or “mismatches” between the two species' thermal performance curves [11,12], can affect inhibitory interactions between symbionts and parasites [13]. This could have important consequences for the temperature dependence of infection. However, few studies have considered the effects of elevated temperature on symbiotic microbiota [14,15], and the consequences of elevated temperature for gut parasite-symbiont competition remain unexplored.

Social bees present an ideal system in which to study effects of temperature on competition between symbionts and parasites. Both honey bees and bumble bees can be infected by a variety of parasites and pathogens, transmission of which is facilitated by the high density of hosts in colonies [16]. However, honey bees and especially bumble bees are facultative endotherms that possess a remarkable ability to regulate their temperatures at the level of the individual bee and colony at over 30 °C above ambient temperature [17,18]. This thermoregulatory capacity allows bumble bees to maintain the temperatures necessary for flight and brood development during times of year when other insects are inactive [19]. The elevated temperatures of bees facilitate not only foraging and colony development, but also defense against infection. In honey bees, high temperatures decreased infection with *Ascosphaera apis* [20], Deformed Wing Virus [21], *Varroa* mites [22], *Nosema apis,* and *N. ceranae* [23].

In addition to their own parasite resistance mechanisms (including thermoregulation), honey and bumble bees have a well-characterized microbiota with demonstrated benefits against infection in larvae and adults [24]. The core gut microbiota consists of five main clades that are found in corbiculate (“pollen-basket”) bees throughout the world [25]. The bumble bee microbiota is dominated by just three of these five core taxa—*Snodgrassella*, *Gilliamella*, and *Lactobacillus* Firmicutes-5 (“Firm-5”)— which together often account for over 80% of the total gut microbiota of worker bumble bees [26–28]. Bacteria isolated from the bumble bee gut had direct inhibitory activity against several bee pathogens [29], and microbiota rich in *Gilliamella* and *Lactobacillus* Firm-5 have been negatively correlated with trypanosomatid infection in bumble bees [26,28,30].

All of the core bumble bee gut symbionts have optimal growth temperatures at 35-37 °C [31,32]. In contrast, widespread trypanosomatid and microsporidian gut parasites (*Crithidia*, *Lotmaria*, and *Nosema* spp.), were described as having optimal temperatures of 25-27 °C [33–35]. This difference in observed *in vitro* growth temperatures suggests the hypothesis that temperatures above the parasites’ thermal optima will favor core symbionts over gut pathogens, due to increased asymmetry in symbiont versus pathogen growth rates at these temperatures. However, no study has empirically quantified differences in the thermal performance curves (i.e., relationship between temperature and growth rate) for symbionts versus parasites, or the temperature dependence of symbiont-mediated parasite inhibition, both of which are likely to shape the effects of temperature on infection in bumble bees.

We used the *Crithidia bombi* / *Lactobacillus bombicola* system to examine temperature dependence of bee symbiont-parasite interactions *in vitro. Crithidia bombi* is an intestinal trypanosomatid that is both widespread and abundant in bumble bees [36,37]. This parasite reduces foraging efficiency and starvation tolerance in worker bees [38,39], growth rates and reproductive output of colonies [40], and post-hibernation survival and hive-founding in queens [38]. Its introduction has been correlated with decline of native bumble bees in South America [41], and its relative *Lotmaria passim* (formerly reported as *C. mellificae*) has been correlated with colony collapse in honey bees [42,43]. *Crithidia bombi* has been cultured at 27 °C [33], and *L. passim* at 25 °C [34]. *Lactobacillus bombicola,* the most widely distributed species found in a cross-species survey of bumble bees [29], is a member of the *Lactobacillus* Firm-5 clade. This clade is found in honey, bumble, and other corbiculate bees worldwide [25]. In honey bees, Firm-5 was the clade with the strongest effect on gut metabolomics [44]. The abundance of Firm-5 bacteria has been negatively correlated with the infection success of *C. bombi* [26,28]. *Lactobacillus bombicola* has been reported to grow at 28-37 °C [45]. Together, these observations suggest that *L. bombicola* is an important gut symbiont that could inhibit *C. bombi* growth in a temperature-dependent manner.

We measured *in vitro* growth of *C. bombi* and *L. bombicola* grown alone, together, and sequentially across a range of incubation temperatures. We tested whether:

1. *Crithidia bombi* and *L. bombicola* growth rates have differential responses to temperature, using metabolic-theory derived models to describe their thermal performance curves,
2. Competitive effects of *L. bombicola* on *C. bombi* increase with temperature and decrease the temperature of peak parasite growth, as predicted based on asymmetries in symbiont versus parasite thermal performance curves, and
3. Temperature-dependent chemical alterations to the growth environment made by *L. bombicola* are sufficient to explain temperature-dependent parasite inhibition.

## MATERIALS AND METHODS

### Overview of experiments

Three series of experiments were conducted to determine the temperature dependence of interactions between *C. bombi* and *L. bombicola*. To estimate Thermal Performance Curves, we measured each species’ growth rate across a range of incubation temperatures. To assess temperature dependence of direct competition, we cocultured *L. bombicola* with *C. bombi* at three incubation temperatures (“Coculture Experiment”). To assess whether a chemical mechanism could explain the temperature-dependent inhibition of parasites in coculture, we compared the effects of *L. bombicola* spent medium from different temperatures on *C. bombi* growth (“Spent Medium Experiment”).

Each experiment used 6 incubators. Thus, for six-temperature experiments used to generate thermal performance curves, we had one incubator-level replicate for each repetition of the experiment. For three-temperature experiments (Coculture and Spent Medium), we had two replicates per repetition. We chose to use the incubator (rather than the sample) as the unit of replication. This accounts for the scale at which the temperature treatment was imposed and avoids pseudoreplication within incubators [3,46]. To increase the number of true replicates of the temperature treatments, we conducted multiple temporal repetitions (blocks) of each experiment, with each incubator assigned to a different temperature treatment during each repetition.

### Cell Cultures

*Crithidia bombi* cell cultures were isolated from bumble bee intestines by flow cytometry-based single cell sorting [33]. Cultures originated from wild infected bumble bees. Strains C1.1 (Corsica, 2009) and S08.1 (Switzerland, 2008, both courtesy Ben Sadd) originated from *B. terrestris*. Strains IL13.2 (Illinois, USA, 2013, courtesy Ben Sadd) and VT1 (Vermont, USA, 2013, courtesy Rebecca Irwin) originated from *B. impatiens*. These same cell lines have been used to assess effects of phytochemicals on parasite growth [47]. Briefly, cells from fecal samples were sorted into 96-well plates containing “FPFB” culture medium with 10% heat-inactivated fetal bovine serum and incubated at 27 °C, then cryopreserved at −80 °C until several weeks before the experiments began [33]. Culture identity was confirmed as *C. bombi* based on glyceraldehyde 3-phosphate dehydrogenase and cytochrome b gene sequences. *Lactobacillus bombicola* strain 70-3, isolated from *Bombus lapidarius* collected near Ghent, Belgium (isolate “28288T” [45]), was obtained from the DSMZ. *Lactobacillus bombicola* was grown in 2 mL screw-cap tubes in MRS broth (Research Products International, Mt. Prospect, IL) with 0.05% cysteine (hereafter “MRSC”) and incubated at 27 °C.

### Thermal performance curves

Growth of each species was measured concurrently by optical density (OD 630 nm) at six temperatures (17-42 °C in 5 °C increments). *Crithidia bombi* cells were added to 96-well plates in 200 μL culture medium at an initial OD of 0.005 (^~^800 cells μL^−1^). OD measurements were taken at 24 h intervals through 120 h of incubation [48]. *Lactobacillus bombicola,* which grew poorly in 96-well plates, was grown in 2 mL screw-cap tubes. Cells were added at an initial OD of 0.020 and measured after 3, 4, 5, 6, and 24 h incubation. The entire experiment was repeated 5 (*C. bombi)* or 6 *(L. bombicola)* times, with incubator temperatures switched between each repetition. Net OD was calculated by subtracting the OD of cell-free medium from the corresponding temperature and time point; this controlled for any changes in OD that occurred independent of cell growth.

We used metabolic theory equations to model the relationship between temperature and growth rate. Growth rates were calculated by fitting a model-free spline [49] to the curve of log-transformed OD (ln(OD_*t*_/OD_t0_)) with respect to time [50]. A separate spline was fit to each replicate combination of incubator, strain, and incubation temperature to estimate the maximum specific growth rate.

Thermal performance curves were modeled for each species and strain using the log-transformed Sharpe-Schoolfield equation [51,52], with temperature as the predictor variable and ln(maximum specific growth rate) as the response variable (Equation 1).

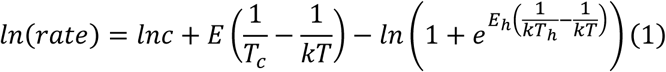

In Equation 1, “*rate*” is the maximum specific growth rate; *lnc* is the natural log of the growth rate at an arbitrary calibration temperature; *E* is the activation energy, which corresponds to the slope of the thermal performance curve below the temperature of peak growth; *T_c_* is the calibration temperature; *k* is Boltzmann’s constant; *T* is the incubation temperature; *E_h_* is the high-temperature deactivation energy, which corresponds to the rate at which growth decreases at supraoptimal temperatures; and T_h_ is the supraoptimal temperature at which growth rate is reduced by 50% relative to peak growth rate.

Solving Equation 1 for the maximum growth rate yields the temperature of peak growth, *T_pk_* [51]:

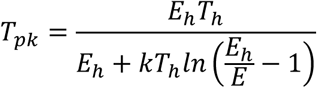

The model fit was optimized for each species and strain using non-linear least squares with package *nls.multstart*, function “nls_multstart” [53]. Model predictions with uncertainty estimates for *T_pk_* and predicted growth at each temperature were estimated by bootstrap resampling (999 iterations). For each bootstrap sample, bootstrap model parameters were estimated, and predictions generated across the full range of incubation temperatures. We constructed 95% bootstrap confidence intervals around the predictions of the original model using the 0.025 and 0.9725 quantiles of predictions from the bootstrap model fits.

### Co-culture Experiment

To assess temperature dependence of direct competition, we cocultured *L. bombicola* with *C. bombi* strain VT1 at three incubation temperatures (27, 32, and 37 °C). These temperatures were chosen for two reasons, one statistical and one physiological. Statistically, these temperatures correspond to the regions of maximal asymmetry in the two species’ thermal performance curves. “Asymmetry” means that the two species have differently or oppositely sloped performance curves across this temperature range [12]. Growth rate of *L. bombicola* continues to increase, whereas growth rate of *C. bombi* plateaus and begins to decline. Physiologically, this is a relevant temperature range for bumble bees. In the hive, thoracic temperatures of workers generally range from 27 to 33 °C (range 23-36 °C), with brood kept near 30 °C [54]. During nest establishment, queens of *Bombus vosnesenskii* maintained even higher temperatures (37.4 to 38.8°C, day and night [54]).

Coculture experiments were conducted in 2 mL tubes in a mixed medium of 50% *Crithidia*-specific FPFB and 50% *Lactobacillus*-specific MRSC. The mixed medium supported growth of both species, whereas neither 100% FPFB (no *L. bombicola* growth) nor 100% MRSC (no *C. bombi* growth) was suitable for coculture. Cells were grown in their preferred treatment media (FPFB for *C. bombi*, MRSC for *L. bombicola)* until the experiment began. At the start of experiment, cells were diluted by OD to 2x final concentrations in their respective preferred media, prior to combination in equal volumes to create the mixed medium. Each experiment included 18 treatments: the three incubation temperatures crossed with two *C. bombi* start densities (initial OD = 0.010 and cell free control) and three *L. bombicola* start densities (initial OD = 0.010, 0.020, and cell free control).

To initiate the experiment, 500 μL each of MRSC-based *L. bombicola* treatment and FPFB-based *C. bombi* treatment were combined in 2 mL screw-cap tubes. Samples were incubated at the appropriate temperature, with growth measurements made after 6 and 24 h of incubation. Growth rates of *L. bombicola* in monoculture were calculated as the rate of increase over the first 6 h (log(OD_6h_/OD_0h_)/6). Growth rates of *C. bombi* in both monoculture and coculture were determined by hemocytometer cell counts at 200x magnification. Quantification of growth by cell counts, rather than OD, allowed us to differentiate growth of the larger, morphologically distinct *C. bombi* from that of *L. bombicola*. Starting cell density was estimated based on cell counts from tubes at time 0 h (OD = 0.010), averaged across all repetitions of the experiment. Final partial OD of *L. bombicola* was approximated by subtracting the estimated OD due to *C. bombi* from the total net OD, using a best-fit linear relationship between *C. bombi* cell density and OD. Growth of *L. bombicola* in coculture was approximated by subtracting the estimated *C. bombi* OD after 6 h of incubation from the total net OD. The calculation assumed constant, exponential growth of *C. bombi* through 24 h incubation, and the same relationship between OD and *C. bombi* cell count observed in the time 0 h samples. Growth rates of *L. bombicola* in coculture should therefore be considered approximate, as we were unable to count individual cells of this species.

Motility of *C. bombi* cells, which are mobile flagellates, was recorded during cell counts. Cell motility of the sample was recorded on an ordinal scale based on whether cells were rapidly swimming, twitching, or immotile (motility scores of 2, 1, and 0, respectively). Monocultured cells were generally the most rapidly moving, and after initial motility screening were diluted in 50% sucrose. The viscosity of this solution slowed the cells to the point where they were countable.

Effects of temperature and *L. bombicola* start density on *C. bombi* growth rate were analyzed by a general linear mixed model [55] with experiment round as a random effect. F-tests were used to evaluate the significance of model terms [56], and pairwise comparisons were made with R package “lsmeans” [57]. Effects of temperature and *L. bombicola* start density on *C. bombi* cell motility were analyzed by a bias-reduced binomial model [58], to cope with complete separation (i.e., no within-group variation in motility). Cell motility was considered as a binary response variable (motility > 0). Likelihood ratio tests were used to evaluate significance of model terms. The relationship between *C. bombi* growth rate and *L. bombicola* OD after 24 h was tested by linear regression.

### Spent medium experiment

We used *L. bombicola* spent medium (i.e., cell-free supernatant of medium in which *L. bombicola* was grown, then removed by filter sterilization) to test whether temperature-dependent inhibition observed in coculture experiments could be explained by temperature-dependent production of inhibitory chemicals by the symbiont. In the first stage of the experiment, *L. bombicola* spent medium was generated at different temperatures. In the second stage, growth of *C. bombi* (strain VT1) was measured in the presence of 50% spent medium at the same temperature at which the spent medium was generated (e.g., spent medium from 32 °C was tested for effects on *C. bombi* incubated at 32 °C; see schematic, Supplementary Figure 1). These experiments used the same three growth temperatures tested in the Coculture Experiment (27, 32, and 37 °C) crossed with three *L. bombicola* start densities (OD of 0, 0.001, or 0.010), for a total of 9 treatments. Each temperature treatment was replicated in two different incubators in each repetition of the experiment. The entire experiment was repeated three times, for a total of six incubator-level replicates.

Spent medium was generated in 14 mL screw-cap conical tubes filled with 8 mL MRSC medium. Each tube was seeded at the appropriate starting density (OD of 0, 0.001, or 0.010) and incubated for 20 h at the appropriate temperature (27, 32, or 37 °C). At the end of the incubation period, a 200 μL aliquot of the resulting spent medium was removed for measurement of OD. The remainder was sterile-filtered to yield the MRSC-based spent medium. A 2 mL aliquot of the spent medium was reserved for measurement of pH; the remainder was used immediately for assays of *C. bombi* growth.

Growth of *C. bombi* was measured in 96-well tissue culture plates in 50% MRSC-based spent medium and 50% *Crithidia*-specific FPFB medium. *Crithidia bombi* cell cultures were diluted to an optical density of 0.020 in *Crithidia*-specific FPFB medium [33]. The *C. bombi* cell suspension (100 μL) was added to an equal volume of spent medium for an initial net OD of 0.010, with 12 replicate wells per plate. Plates were incubated at the same temperature used for generation of the spent medium. Growth was measured by OD at 20, 26, 44, and 50 h of incubation. Net OD was computed by subtraction of OD from cell-free control wells of the corresponding spent medium treatment and time point. Visual inspection of growth curves indicated that maximum growth rate occurred during the initial incubation interval (0-20 h). Therefore, relative growth rate was computed as

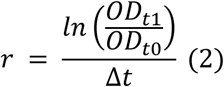

Where OD_t0_ represents initial OD of *C. bombi* (0.010), OD_t1_ represents OD at the time of first measurement (20 h), and Δt is the amount of time between the start of the experiment and the first measurement.

Effects of temperature and *L. bombicola* start density on *C. bombi* growth rate were analyzed by a general linear mixed model with experiment round as a random effect [55]. F-tests were used to evaluate the significance of model terms [56]. Relationships between spent medium OD before filtration and temperature, spent medium pH and temperature, *C. bombi* growth rate and spent medium OD before filtration, and *C. bombi* growth rate and spent medium pH were tested by linear regression.

## RESULTS

**Thermal performance curves showed higher temperatures of peak growth and upper limits of thermotolerance in *L. bombicola* than in *C. bombi* (Fig 1).** All *C. bombi* strains showed similar model-predicted peak growth temperatures (T_pk_), ranging from 33.9 °C in strain S08.1 to 34.4 °C in strain IL13.2. These estimates were at least 5 °C lower than the estimated T_pk_ for *L. bombicola* (39.83 °C, Fig. 1). For all strains of *C. bombi,* the temperature that inhibited growth by 50% (T_h_) was below 38 °C, or at least 5 °C lower than the T_h_ for *L. bombicola* (Fig. 1). Full model parameters are given in Supplementary Table S1.

**Figure 1.**
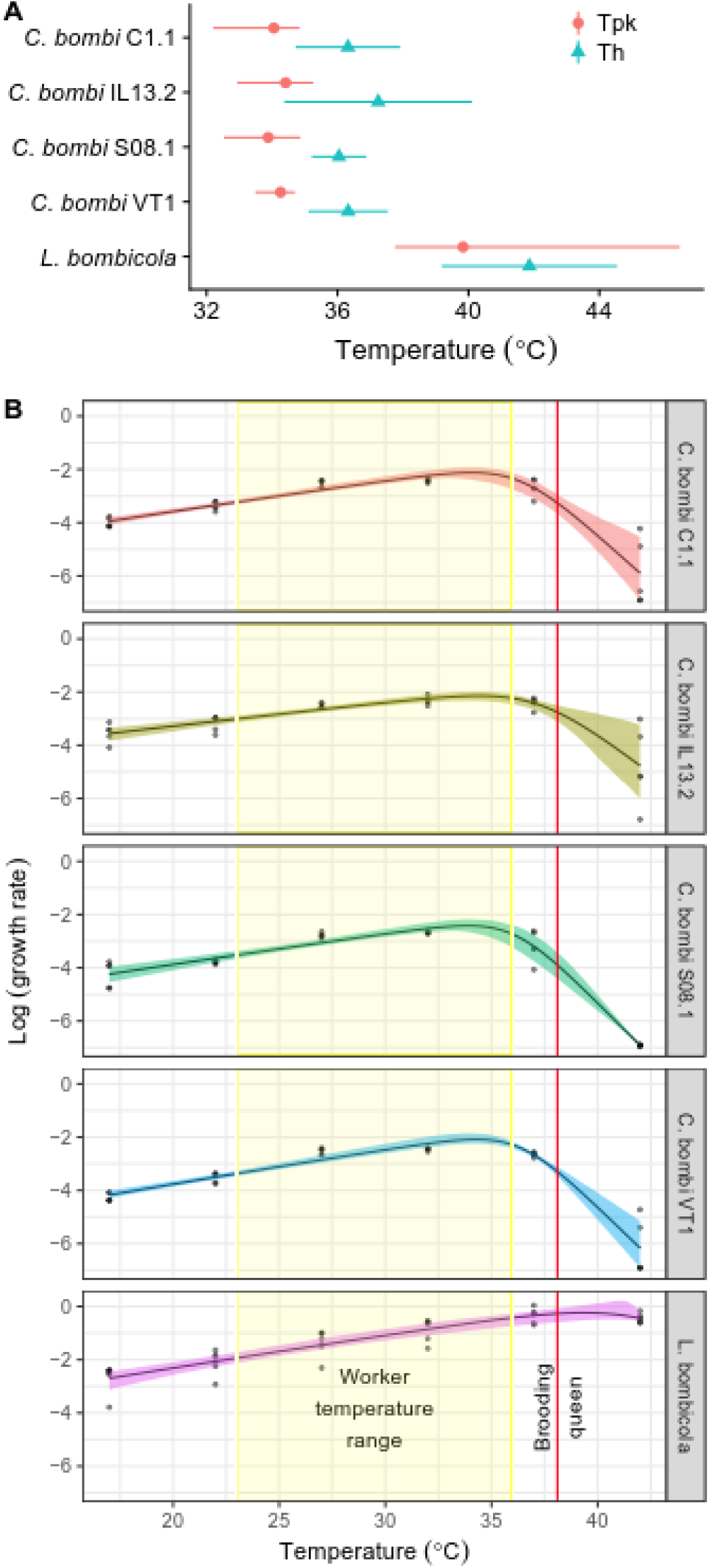
*L. bombicola* exhibited higher peak growth temperature and greater tolerance to high temperatures than did *C. bombi*. **(A)** Model parameters for four *C. bombi* strains and *L. bombicola*. Points and error bars show means and 95% confidence intervals for peak growth temperature (Tpk, based on predictions from Sharpe-Schoolfield model from 999 bootstrap samples) and temperature at which growth was reduced by 50% relative to peak growth (Th, based on Sharpe-Schoolfield model fit by nonlinear least squares). **(B)** Full thermal performance curves used to derive model parameters shown in (A). Y-axis shows log-transformed specific growth rate (μ (h-1)) based on spline fits. Points show raw data, with one point per replicate (incubator). Trendlines show predictions from Sharpe-Schoolfield models. Shaded bands show 95% bootstrap confidence intervals. The curves are overlain with physiologically relevant temperature ranges for bumble bee workers (yellow vertical region) and queens (red vertical line), using data from [54]. Please refer to online version of article for color figure.

**Coculture with *L. bombicola* inhibited *C. bombi* growth and motility, and reduced temperature of peak *C. bombi* growth (Fig. 2).** Growth rate of *C. bombi* was reduced by over 50% in coculture (temperature-adjusted marginal mean 0.66 ± 0.005 in monoculture vs. 0.32 ± 0.005 in coculture, Fig. 2A). We found stronger inhibitory effects of *L. bombicola* at higher temperatures (temperature x *L. bombicola* start density interaction, F_4, 43_ = 3.30, P = 0.019). Competition with *L. bombicola* altered the shape of the *C. bombi* thermal performance curve. Whereas *C. bombi* grew well throughout the range of 27-37 °C in monoculture, growth was poor above 27 °C in coculture (Fig. 2A). In addition to reducing growth, coculture with *L. bombicola* profoundly reduced *C. bombi* cell motility in a temperature-dependent fashion (Temperature x *L. bombicola* interaction: Chi-squared = 16.36, Df = 1, P < 0.001, Fig. 2B). Whereas cells remained motile regardless of temperature in monoculture, no motility was observed above 27 °C in coculture. The stronger effects of *L. bombicola* on *C. bombi* at high temperatures reflected increased *L. bombicola* cell densities, which were negatively correlated with *C. bombi* growth rate (estimate = −0.094 ± 0.013 SE, t = −7.52, P < 0.001, R^2^ = 0.521).

**Figure 2.**
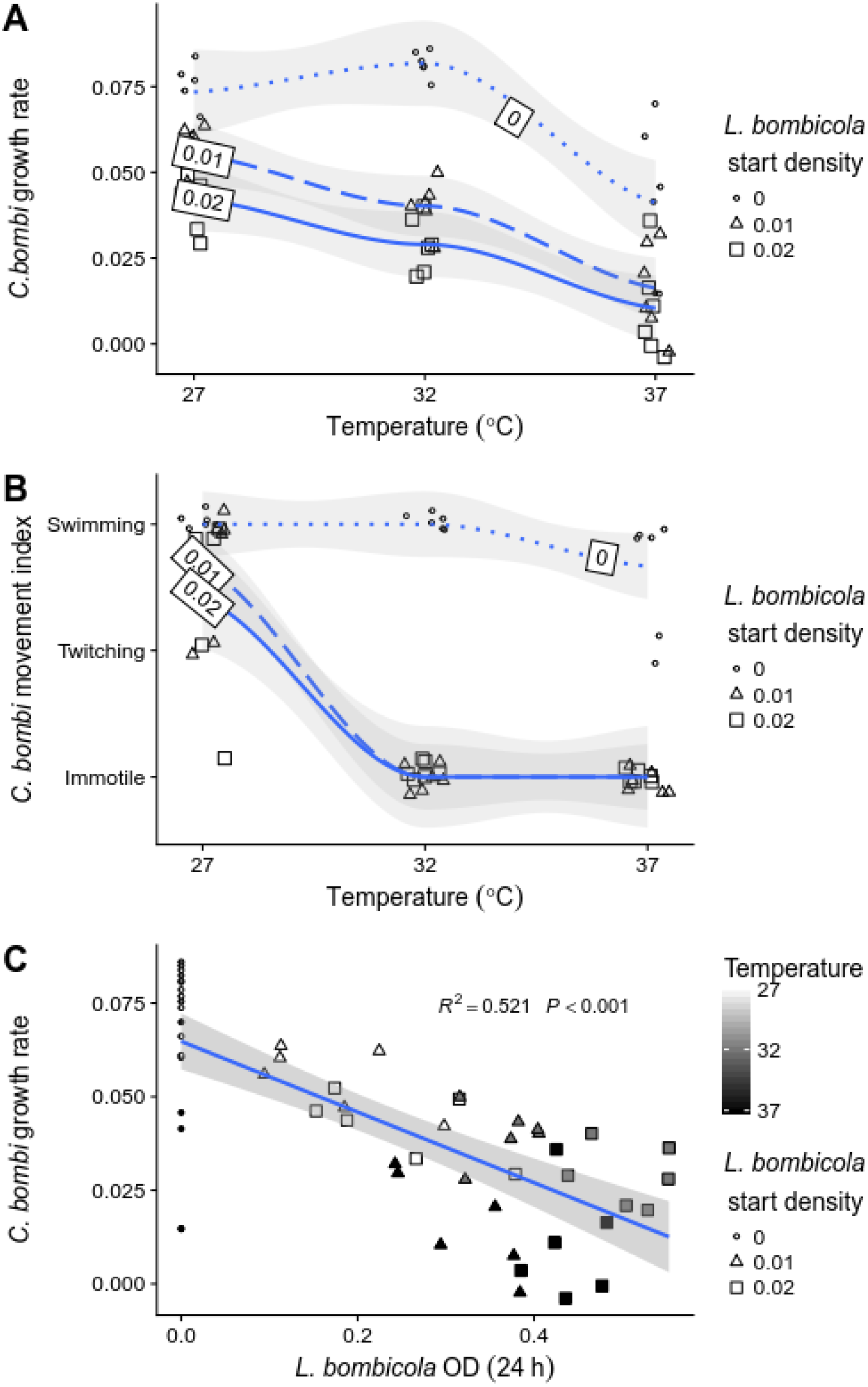
Competition with *L. bombicola* inhibited growth of *C. bombi* and reduced peak growth temperature, due to higher *L. bombicola* growth rates at high temperatures. **(A)** *Crithidia bombi* growth rate at 3 different temperatures in the presence of 3 starting optical densities (OD) of *L. bombicola*: 0 (i.e., no *L. bombicola*, small hollow circles and dotted line), 0.01 (medium-sized triangles and dashed line), and 0.02 (large squares and solid line). Each point represents specific growth rate (μ (h-1)) based on cell counts for a single incubator and repetition of the experiment. Trendlines show smoothed Loess fits for each *L. bombicola* start density; shaded bands show 95% confidence intervals. Points have been randomly offset to reduce overplotting. **(B)** *Crithidia bombi* cell motility, observed microscopically after 24 h of coculture at the time of cell counts used to calculate growth rates in (A). Points have been randomly offset to reduce overplotting. Symbol size, symbol shape, and trendlines match interpretations for panel (A). No movement of *C. bombi* was observed for any of the *C. bombi* cocultured with *L. bombicola* at 32 or 37°C. **(C)** *Crithidia bombi* growth rate was negatively correlated with OD of *L. bombicola* after 24 h of coculture. Partial OD of *L. bombicola* was estimated as net OD after subtraction of estimated OD due to *C. bombi,* based on correlation between OD and *C. bombi* cell concentration. Symbol fill indicates temperature; symbol shape and size indicate *L. bombicola* start density. Trendline shows linear model fit; shaded band shows 95% confidence interval.

Whereas *L. bombicola* had negative effects on *C. bombi, C. bombi* appeared to increase growth rate of *L. bombicola* under the conditions of our experiments. Estimated *L. bombicola* growth rate was nearly 3-fold higher in the presence of *C. bombi* than in its absence (temperature-adjusted mean growth rate = 0.515 ± 0.008 SE with *C. bombi* vs. 0.181 ± 0.008 SE without *C. bombi,* t = 29.8, P < 0.001, Supplementary Figure 2).

***Lactobacillus bombicola* spent medium reduced *C. bombi* growth rate and peak growth temperature (Fig. 3).** As in the Coculture Experiment, we found temperature-dependent inhibition of *C. bombi* by *L. bombicola* in the Spent Medium Experiment. Whereas *C. bombi* grew well at all temperatures in control medium, growth was decreased at high temperatures in the presence of *L. bombicola* spent medium produced at high temperatures (Temperature x *L. bombicola* start density interaction: F_4, 43_ = 8.28, P < 0.001, Fig. 3A). The stronger inhibitory effects of spent medium from higher temperatures reflected faster growth of *L. bombicola* at higher temperatures, which led to greater OD (t = 3.56, P < 0.001) and lower pH (t = −3.84, P < 0.001) achieved at higher temperatures during generation of the spent medium. As in the Coculture Experiment, *C. bombi* growth rate was negatively correlated with final OD of *L. bombicola* (estimate = −0.083 ± 0.017 SE, t = −4.79, P < 0.001, R^2^ = 0.29, Fig. 3B), and even more strongly negatively correlated with acidity of spent medium (effect of pH: estimate = 10.72 ± 1.64, t = 6.54, P < 0.001, R^2^ = 0.44, Fig. 3C).

**Figure 3.**
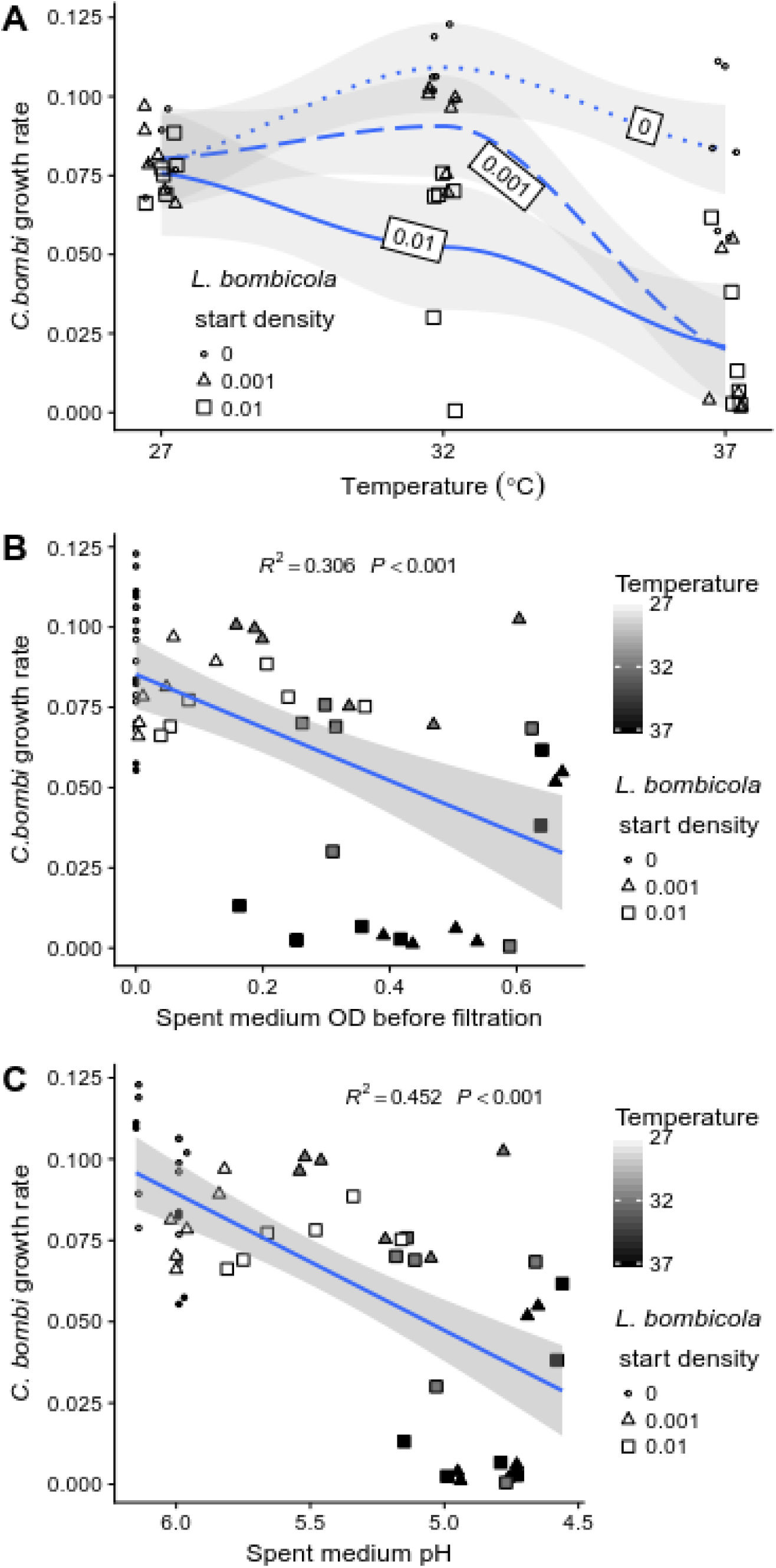
Spent medium from *L. bombicola* reduced growth rate and peak growth temperature of *C. bombi*, due to higher rates of *L. bombicola* growth and acid production at high temperatures. **(A)** *Crithidia bombi* growth rate at 3 different temperatures in the presence of spent medium. Spent medium was generated by growth of *L. bombicola* for 24 h from 3 starting densities: OD = 0 (i.e., no *L. bombicola*, small hollow circles and dotted line), 0.001 (medium-sized gray circles and dashed line), and 0.01 (large black circles and solid line). Each point represents specific growth rate (μ (h-1)) based on cell counts for a single incubator and repetition of the experiment. Trendlines show smoothed Loess fits; shaded bands show 95% confidence intervals. **(B)** *Crithidia bomb*i growth rate was negatively correlated with OD of *L. bombicola* at the time when spent medium was filtered (i.e., after 24 h incubation). Symbol fill indicates temperature; symbol shape and size indicate *L. bombicola* start density. Note higher OD’s achieved at higher temperatures, except in the *L. bombicola*-free controls (start density = 0, circles). Growth of *C. bombi* was assayed at the same temperature at which the spent medium had been generated. Trendline shows linear model fit, pooled across start densities and temperatures. Shaded band shows 95% confidence interval. **(C)** Growth rate of *C. bombi* was negatively correlated with acidity of *L. bombicola* spent medium. X-axis shows pH of spent medium after 20 h growth of *L. bombicola,* at the beginning of *the C. bombi* growth assay. As in (B), symbol fill indicates incubation temperature, symbol shape and size indicate *L. bombicola* start density, and trendline with shaded band shows linear model fit with 95% confidence bands. Note higher acidity (lower pH) achieved at higher temperatures, except in the *L. bombicola*-free controls (start density = 0, circles).

## DISCUSSION

As expected based on temperatures conventionally used in cell cultures, the symbiont *L. bombicola* had higher temperatures of peak growth and grew at higher temperatures than those tolerated by the parasite *C. bombi*. All four tested parasite strains exhibited similar thermal performance curves and inhibitory temperatures. This was somewhat surprising given the documented among-strain variation in growth rate [59], infectivity [60], and ability to tolerate dessication [61], phytochemicals [48], antimicrobial peptides [62], and gut microbiota [26]. The conservation of thermal performance profiles across strains could reflect strong stabilizing selection for enzymes and metabolic processes involved in thermotolerance, or adaptation to a consistent range of temperatures experienced in the bee abdomen. Regardless of the physiological underpinning, consistent upper limits of thermotolerance across parasite strains suggest that elevated temperature would be an effective defense against a range of *C. bombi* parasite genotypes.

The differently shaped thermal performance curves of *L. bombicola* and *C. bombi* indicate that a temperature increase over the range recorded in bumble bees would favor growth of symbionts over parasites, while the inhibitory effects of *L. bombicola* on *C. bombi* indicate that this increased symbiont growth could constrain the ability of parasites to persist at high temperatures. Growth rates of *C. bombi* plateaued over the 27-33 °C range found in bumble bee nests [54], and began to drop at the 38 °C temperatures found in post-hibernation queens [54], the life stage at which bumble bees are most vulnerable to the effects of *C. bombi* [38]. In contrast, growth rate of *L. bombicola* continued to increase throughout this interval, rising nearly three-fold from 0.265 h^−1^ at 27 °C to 0.734 h^−1^ at 37 °C. As a result, any effects of *L. bombicola* on *C. bombi* should become more pronounced at higher temperatures.

Within the gut, interactions between species may be positive, negative, or neutral. For example, the bee gut symbionts *Snodgrassella* and *Gilliamella* facilitate one another’s growth physically, via formation of multi-species biofilms [63], and chemically, via cross-feeding and modification of gut oxygen concentration and pH [44,64]. The effects of *L. bombicola* on *C. bombi* were strongly inhibitory. We have shown this inhibition to be chemically mediated by *L. bombicola*’s production of acids [65]. Because *L. bombicola* rates of growth and acid production increased over the temperature range found in bees, we predict that increases in bee body temperature would reduce infection by increasing growth rate of *L. bombicola* and related Firm-5 bacteria, thereby decreasing gut pH to the point where parasites cannot grow. Thus, although parasites in monoculture are capable of growth throughout the range of temperatures found in bees, our results predict that competitive exclusion by symbionts could limit the parasite’s thermal niche to cooler temperatures.

In contrast to the inhibitory effects of *L. bombicola* on *C. bombi, C. bombi* appeared to facilitate growth of *L. bombicola*. Given that *L. bombicola* did not grow at all in full-strength FPFB medium, this facilitation could reflect *C. bombi*’s catabolism of *L. bombicola-inhibitory* components, such as serum, in the mixed MRSC/FPFB growth medium. Still, our findings indicate highly asymmetric competition between these two species, to the advantage of the symbiont.

The equilibrium outcome of competitive interactions depends on both interaction strengths and initial densities [66]. In the case of *L. bombicola* and *C. bombi,* initial symbiont densities had the strongest effects at intermediate temperatures typical of a bumble bee hive (27-33 °C). At these moderate temperatures, lower symbiont and higher parasite growth rates might allow parasites to establish if initial symbiont densities are low. In contrast, at higher temperatures typical of those found in queens (>37 °C), high symbiont growth rates and direct high-temperature inhibition of parasites quickly made up for low initial symbiont density. In the social *Bombus* and *Apis* bees, core symbionts such as *Lactobacillus* Firm-5 are rapidly acquired by newly emerged bees from nestmates and hive materials [27,67]. This socially mediated inoculation with core symbionts can establish a protective barrier against infection in colonies with microbiota that contain acid-producing *Gilliamella* and *Lactobacillus* Firm-5 [26,28]. However, symbiont-based defenses might be weakened by treatment with antibiotics, which reduced populations of core gut symbionts and resistance to *C. bombi* [68]. Symbiont-based defenses might also be relatively weak in solitary bees, which can be infected by the same trypanosomatids that infect honey and bumble bees [34]. These bees lack a thermoregulated nest environment and a socially transmitted core gut microbiota, instead acquiring acidophilic gut symbionts from their environment [69]. As a result, solitary bees might be vulnerable to trypanosomatid infection during maturation of their gut microbiota, especially at cooler temperatures. However, no study has experimentally investigated trypanosomatid infections in solitary bee species, let alone the temperature dependence of such infection.

In our *in vitro* host-parasite-symbiont system, we found that high temperatures favored symbionts over pathogens. This suggests that infection-related increases in body temperature, such as fever, may allow hosts to clear pathogens while sparing beneficial symbionts. However, maintenance of elevated temperature comes at an energetic cost in both endothermic mammals and insects such as bumble bees [17,54]. In bees and other endothermic hosts, the ability to maintain parasite-inhibiting temperature will depend on sufficient caloric resources. Further study of temperature-dependent changes to microbiota and infection in live bees, and the effects of infection on endogenous thermoregulation and temperature preference, will be necessary to determine how our *in vitro* findings scale up to the organismal scale.

Studies of other host-symbiont-parasite systems are needed to determine whether high temperatures achieved during febrile states can be detrimental to symbiont populations [14], whether directly or via upregulation of host immunity [8,70], and the consequences of these effects for infection and host health. For example, short-term heat exposure altered soil microbial communities, and caused
loss of the soil’s activity against plant diseased [71]. Numerous examples demonstrate that depletion of symbionts increases susceptibility to infection in animals as well [68,72–74]. Amidst growing appreciation for the roles of temperature, fever, and the microbiome in infectious disease, understanding the effects of temperature on microbiota-parasite interactions may help to predict infection outcome in animals that exhibit fever, and in ectotherms that face infection in changing climates.

## ACKNOWLEDGMENTS

The authors thank Ben Sadd for providing strains C1.1, IL13.2 and S08.1, Rebecca Irwin for providing the bees from which Strain VT1 was established, the DSMZ for providing *L. bombicola,* Guang Xu and Ben Sadd for sharing DNA sequences, and Daniel Padfield for sharing R script.

## DATA AVAILABILITY

All data are supplied in the Supplementary Information, Data S1.

## AUTHORS’ CONTRIBUTIONS

ECPY and QSM conceived the study. ECPY designed and conducted the experiments and analyzed the data with guidance from TRR and QSM. ECPY drafted the manuscript. All authors revised the manuscript and gave final approval for publication.

## MEDIA PROMOTION

Many animals use elevated body temperature (fever) and beneficial gut bacteria to combat infection. However, effects of high temperatures on competition between beneficial and pathogenic microbes remain unknown. We tested effects of temperature on competition between a gut pathogen and a symbiont of bumble bees—insects threatened by disease, but capable of elevating their nest and body temperatures. An increase in temperature over the range found in bee colonies favored beneficial bacteria over pathogens. This suggests that high body temperatures might reduce infection by clearing pathogens while sparing beneficial bacteria, highlighting an unexplored mechanism by which fevers could ameliorate disease.

## SUPPLEMENTARY FIGURES

**Supplementary Figure 1.**
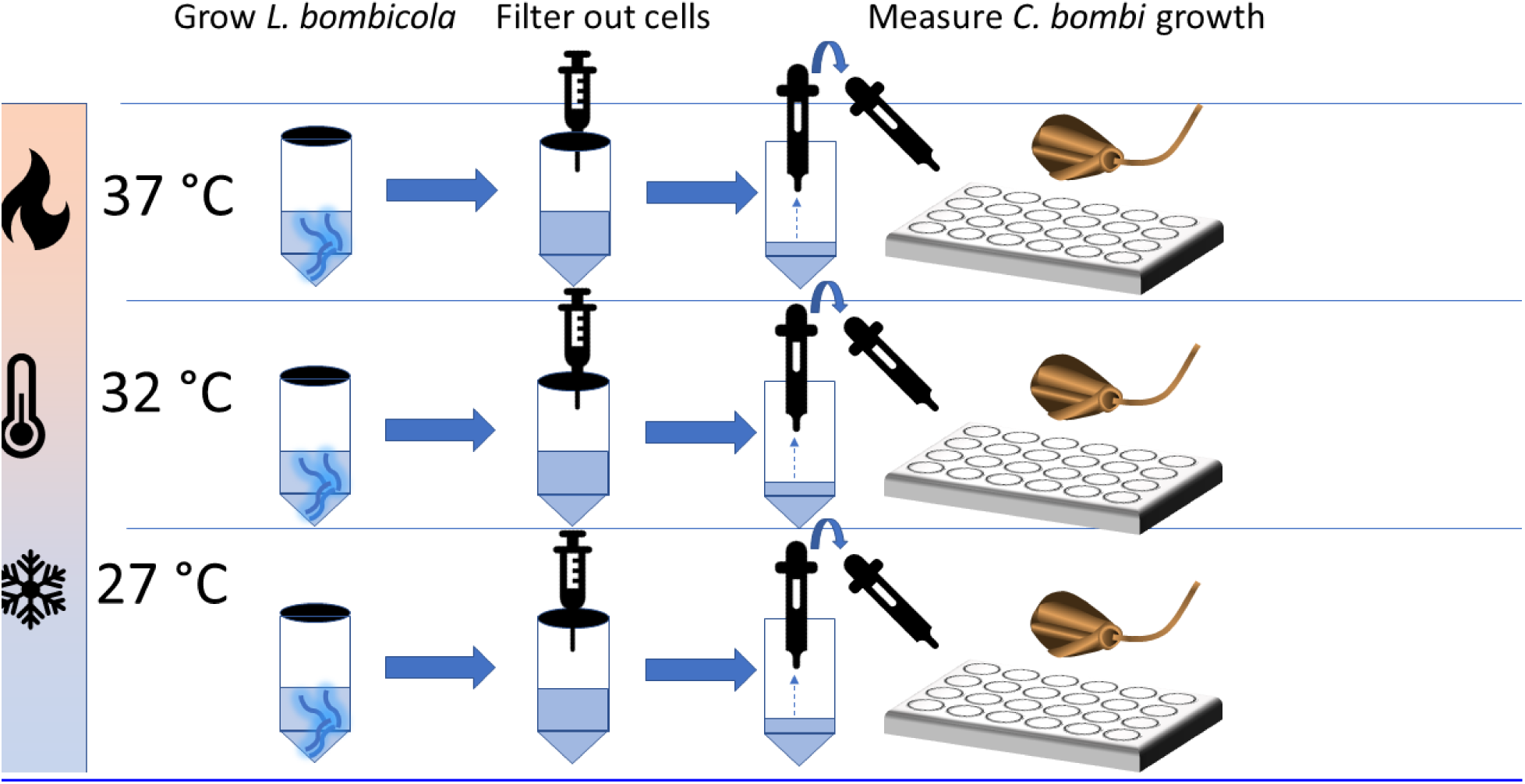
Schematic of Spent Medium Experiment. To generate spent medium, *L. bombicola* was grown for 20 h in screw-cap conical tubes at one of three temperatures (27, 32, or 37 °C). At the end of the incubation period, the resulting medium was sterile-filtered to yield the MRSC-based spent medium, which was used immediately for assays of *C. bombi*growth. The temperature of the *C. bombi* growth assay matched the temperature at which the spent medium had been generated. The entire experiment was repeated three times, each time using two incubators for each of the three temperatures.

**Supplementary Figure 2.**
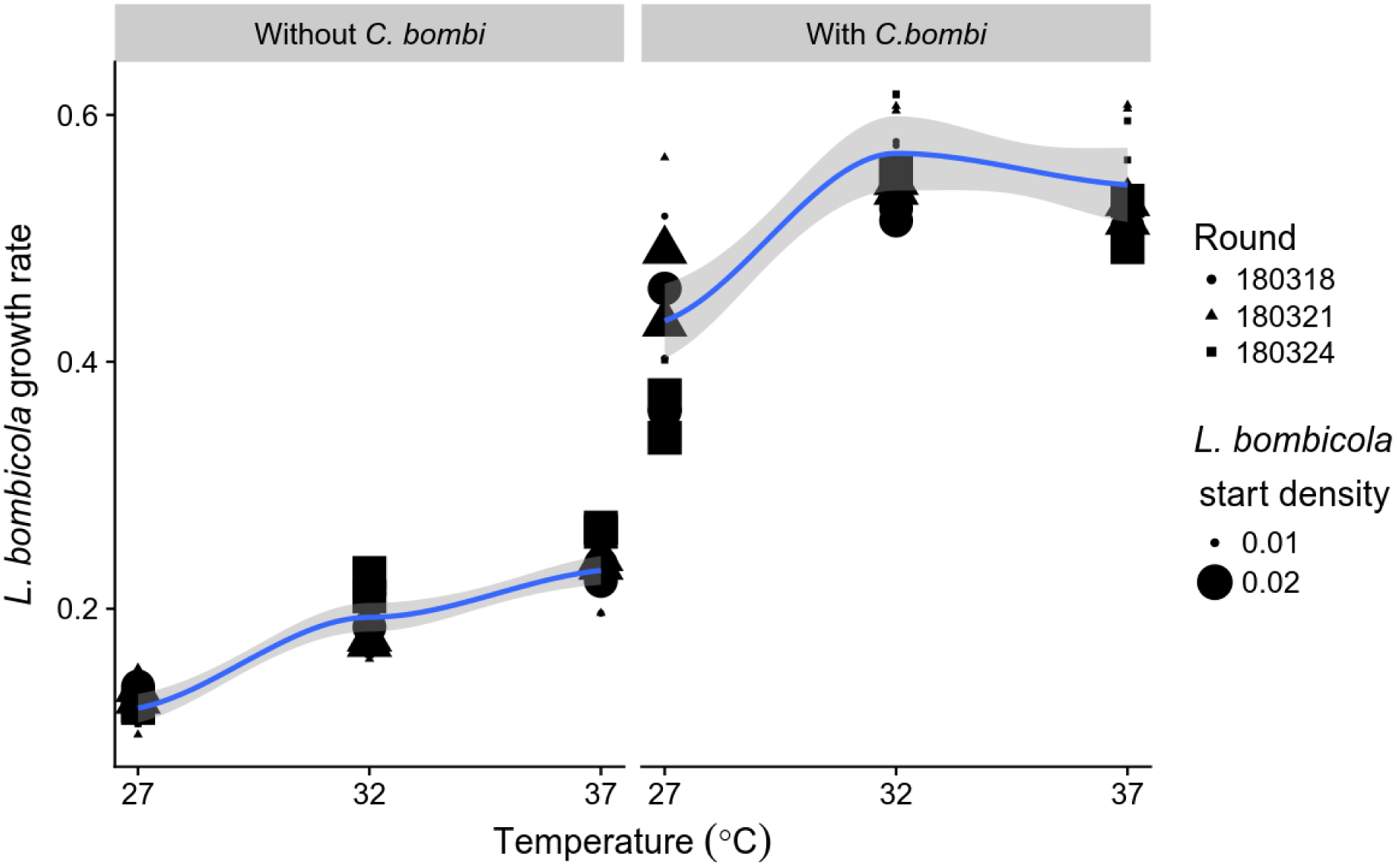
Estimated *L. bombicola* growth rates were higher in the presence of *C. bombi* than in its absence. Left panel, *L. bombicola* monocultures without *C. bombi*. Right panel, cocultures with *C. bombi* start density of OD = 0.010. Shapes represent different repetitions of the experiment, each with 2 incubators per incubation temperature. Symbol size represents *L. bombicola* start density.

## SUPPLEMENTARY TABLES

**Supplementary Table 1.** Parameter estimates and confidence intervals for thermal performance curves. Tpk was estimated by bootstrapping, and has no associated standard error because it is not an explicit model parameter. Temperatures (Th and Tpk) are given in Kelvin.

| Strain | term | estimate | std.error | conf.low | conf.high |
| --- | --- | --- | --- | --- | --- |
| *L. bombicola* | Inc | -2.314 | 0.105 | -2.528 | -2.100 |
| *L. bombicola* | E | 0.941 | 0.123 | 0.690 | 1.191 |
| *L. bombicola* | Eh | 4.645 | 6.866 | -9.340 | 18.630 |
| *L. bombicola* | Th | 315.015 | 1.316 | 312.334 | 317.696 |
| *C. bombi* C1.1 | Inc | -3.585 | 0.154 | -3.901 | -3.269 |
| *C. bombi* C1.1 | E | 0.881 | 0.169 | 0.533 | 1.230 |
| *C. bombi* C1.1 | Eh | 7.014 | 0.774 | 5.424 | 8.604 |
| *C. bombi* C1.1 | Th | 309.469 | 0.778 | 307.870 | 311.068 |
| *C. bombi* IL13.2 | Inc | -3.284 | 0.177 | -3.649 | -2.920 |
| *C. bombi* IL13.2 | E | 0.680 | 0.199 | 0.271 | 1.090 |
| *C. bombi* IL13.2 | Eh | 5.897 | 1.275 | 3.276 | 8.518 |
| *C. bombi* IL13.2 | Th | 310.394 | 1.395 | 307.527 | 313.261 |
| *C. bombi* S08.1 | Inc | -3.877 | 0.097 | -4.076 | -3.678 |
| *C. bombi* S08.1 | E | 0.883 | 0.106 | 0.665 | 1.100 |
| *C. bombi* S08.1 | Eh | 7.718 | 0.447 | 6.799 | 8.636 |
| *C. bombi* S08.1 | Th | 309.195 | 0.405 | 308.362 | 310.027 |
| *C. bombi* VT1 | Inc | -3.766 | 0.131 | -4.035 | -3.498 |
| *C. bombi* VT1 | E | 0.991 | 0.142 | 0.698 | 1.284 |
| *C. bombi* VT1 | Eh | 7.602 | 0.646 | 6.274 | 8.930 |
| *C. bombi* VT1 | Th | 309.475 | 0.590 | 308.262 | 310.689 |
| *C. bombi* C1.1 | Tpk | 307.202 | N/A | 305.351 | 307.991 |
| *C. bombi* IL13.2 | Tpk | 307.564 | N/A | 306.082 | 308.415 |
| *C. bombi* S08.1 | Tpk | 307.031 | N/A | 305.677 | 308.007 |
| *C. bombi* VT1 | Tpk | 307.408 | N/A | 306.637 | 307.850 |
| *L. bombicola* | Tpk | 312.986 | N/A | 310.907 | 319.610 |

